# ClusterMap: multi-scale clustering analysis of spatial gene expression

**DOI:** 10.1101/2021.02.18.431337

**Authors:** Yichun He, Xin Tang, Jiahao Huang, Haowen Zhou, Kevin Chen, Albert Liu, Jingyi Ren, Hailing Shi, Zuwan Lin, Qiang Li, Abhishek Aditham, Jian Shu, Jia Liu, Xiao Wang

## Abstract

Quantifying RNAs in their spatial context is crucial to understanding gene expression and regulation in complex tissues. *In situ* transcriptomic methods generate spatially resolved RNA profiles in intact tissues. However, there is a lack of a unified computational framework for integrative analysis of *in situ* transcriptomic data. Here, we present an unsupervised and annotation-free framework, termed ClusterMap, which incorporates physical proximity and gene identity of RNAs, formulates the task as a point pattern analysis problem, and thus defines biologically meaningful structures and groups. Specifically, ClusterMap precisely clusters RNAs into subcellular structures, cell bodies, and tissue regions in both two- and three-dimensional space, and consistently performs on diverse tissue types, including mouse brain, placenta, gut, and human cardiac organoids. We demonstrate ClusterMap to be broadly applicable to various *in situ* transcriptomic measurements to uncover gene expression patterns, cell-cell interactions, and tissue organization principles from high-dimensional transcriptomic images.

---

Tissue functions arise from the orchestrated interactions of multiple cell types, which are shaped by differential gene expression in three-dimensional (3D) space. To chart the spatial heterogeneity of gene expression in cells and tissues, a myriad of image-based *in situ* transcriptomics methods (*e*.*g*., STARmap, FISSEQ, pciSeq, MERFISH, seqFISH, osmFISH, *etc*.) have been developed^1-8^, providing an atlas of subcellular RNA localization in intact tissues. However, it is challenging to directly extract low-dimensional representations of biological patterns from high-dimensional spatial transcriptomic data.

One main challenge is to achieve accurate and automatic cell segmentation that accurately assigns RNAs into individual cells for single-cell analysis. The most common cell segmentation strategy is labeling cell nuclei or cell bodies by fluorescent staining^9-11^ (*e*.*g*., DAPI, Nissl, WGA, *etc*.) and then segmenting the continuous fluorescent signals by conventional or machine learning (ML)-based methods^12^. However, conventional methods, such as distance-transformed watershed^13^, require manual curation to achieve optimal segmentation. On the other hand, while ML-based methods^14,15^ can automatically detect the targets (cells) in fluorescent stainings, they still require manually annotated datasets for model training and have poor generalization ability to other datasets.

In order to address these challenges, a fundamentally different approach that bypasses auxiliary cell staining, hyperparameter tuning, and manual labeling is warranted. Here, instead of using fluorescent staining, we directly utilized the patterns of spatially resolved RNAs that intrinsically encode high-dimensional gene expression information for subcellular and cellular segmentation, followed by cell-type spatial mapping. To leverage the spatial heterogeneity of RNA-defined cell types, we applied the same strategy to cluster discrete cells into tissue regions. Together, we demonstrated that this computational framework (termed ClusterMap) can identify subcellular structures, cells, and tissue regions (Fig. 1).

**Fig. 1:**
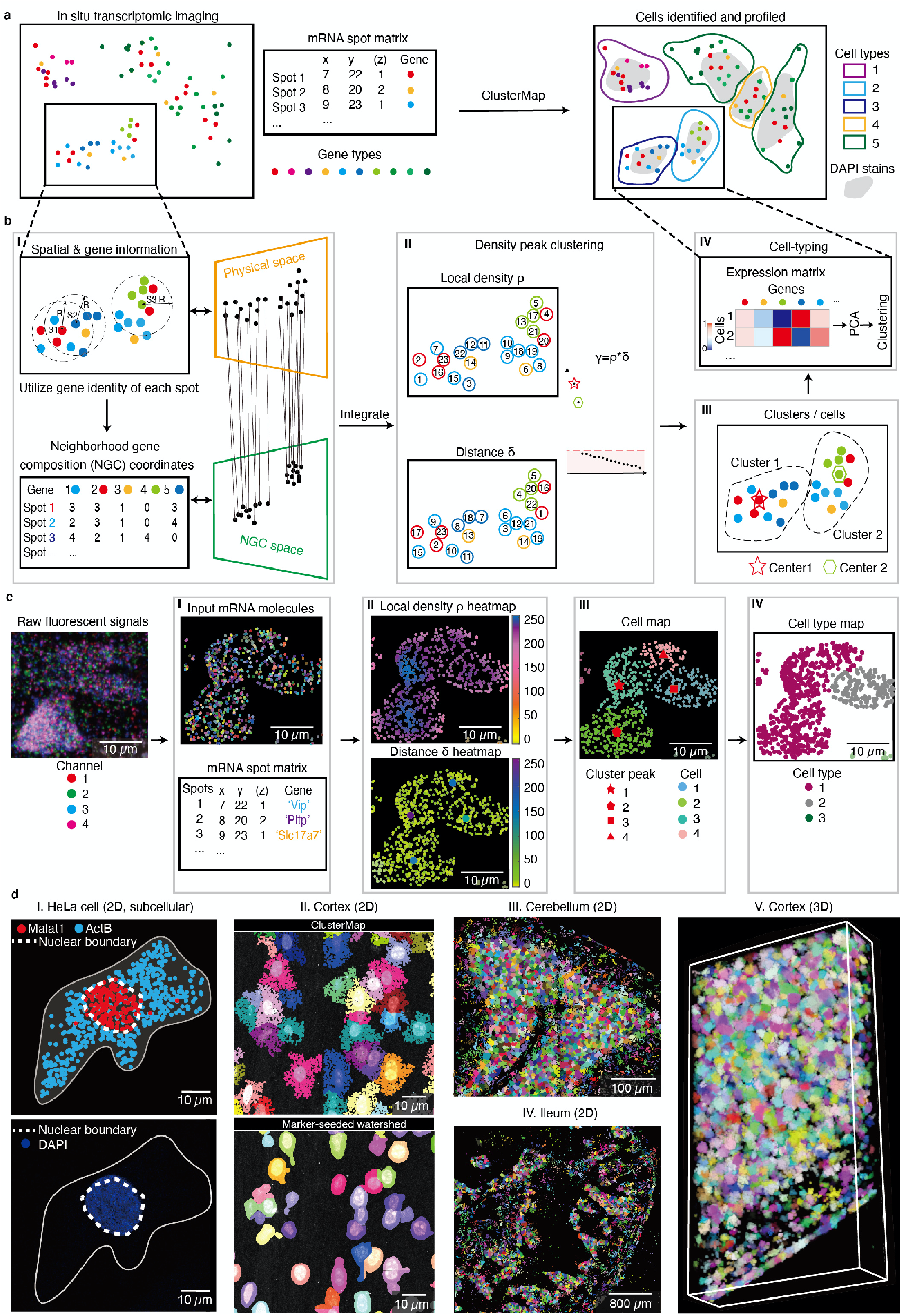
ClusterMap: multi-scale spatial clustering analysis of *in situ* transcriptomic data from subcellular to tissue scales. **a**, Overview of ClusterMap method. The input is a matrix that contains both spatial and transcript information of mRNA molecules sequenced by *in situ* transcriptomic methods^1-8^. ClusterMap clusters mRNA spots, identifies cells, and profiles them into different cell types as output. **b**, Workflow of ClusterMap method. **I**, The physical and neighborhood gene composition (NGC) coordinates of mRNA spots are extracted for each spot (*e*.*g*., S1, S2, and S3), and projected to physical and NGC spaces respectively, which are then computationally integrated. **II**, Density peak clustering (DPC) algorithm^18^ is used to cluster mRNA in the P-NGC space. **III**, Each spot is assigned to one cluster, representing one cell. **IV**, Cell types are identified by the gene expression profiles in each cell. **c**, Representative ClusterMap analysis on STARmap mouse V1 1020-gene dataset^6^ corresponds to (**I**)-(**IV**) in (**b**). **d**, Representative ClusterMap cell segmentation analysis on different samples. **I**, HeLa cell in 2D. The white dashed lines highlight the nuclear boundary identified by the subcellular mRNA distribution from ClusterMap (upper) and DAPI staining (bottom) from the same cell. **II**. Comparison of ClusterMap (upper) and marker-seeded watershed (bottom) segmentation in mouse visual cortex cells. **III**, Mouse cerebellum in 2D, 4,050 cells. **IV**, Mouse ileum in 2D, 5,550 cells. **V**, Mouse visual cortex in 3D, 2,251 cells. Width: 309 µm, height: 582 µm, depth: 100 µm.

## Results

### ClusterMap integrates spatial and gene expression analyses

ClusterMap is based on two key biological phenomena. First, the density of RNA molecules is higher inside cells than outside cells; second, cellular RNAs encoded by different genes are enriched at different subcellular locations, cell types, and tissue regions^16,17^. Thus, we reasoned that we could identify biologically meaningful patterns and structures directly from *in situ* transcriptomic data by joint clustering the physical density and gene identity of RNAs. Subsequently, the spatial clusters were interpreted based on the gene identity and spatial scales to represent subcellular localization, cell segmentation, and region identification. ClusterMap started with pre-processed imaging-based *in situ* transcriptomic data (Methods), where raw fluorescent images were converted into discrete RNA spots with a physical 3D location and a gene identity (*i*.*e*. mRNA spot matrix, Fig. 1a). We reasoned that spatial clusters can be distinguished based on the gene expression in the local neighborhood of each RNA spot. To quantify this, we introduced multidimensional coordinates, termed neighborhood gene composition (NGC) coordinates, which were computed by considering gene expression profiles in a circular window over each RNA spot (Fig. 1b (I), Methods). ClusterMap is capable of analysis on different spatial resolutions by changing the size of the window. The size of the window is specifically chosen for the same dataset to match the average size of organelles or cells for subcellular or single-cell analysis, respectively (Methods). The NGC coordinates and physical coordinates of each RNA spot are then computationally integrated into joint physical and NGC (P-NGC) coordinates over each spot.

Next, we aimed to cluster the RNAs in the P-NGC coordinates for downstream segmentation. Out of numerous clustering algorithms, density peak clustering (DPC)^18^, a type of density-based clustering method, was chosen for its versatility in extracting biological features in data and its compatibility with clusters of various shapes and dimensionalities automatically. DPC identifies cluster centers with a higher density than the surrounding regions as well as a relatively large distance from points with higher densities. We applied DPC to compute two variables^18^: local density *ρ* and distance *δ* for each spot in the joint P-NGC space. For each spot, *ρ* value represents the density of its closely surrounded spots, and *δ* value represents the minimal distance to spots with higher *ρ* values. Spots with both high *ρ* and *δ* values are highly likely to be cluster centers. We then ranked the product of these two variables, *γ*, in decreasing order to find genuine clusters with orders of magnitude higher *γ* values (Methods). For example, in Fig. 1b, the two spots with the *γ* values that are orders of magnitude higher than other spots are chosen as cell centers (labeled by a red star and a cyan hexagon, Fig. 1b (II)). After selecting the two cluster centers, the remaining spots are assigned to one of the clusters respectively in a descending order of *ρ* value. Each is assigned to the same cluster as its nearest cluster-assigned neighbor^14^, and each cluster of spots is taken to represent an individual cell (Fig. 1b (III)). Outliers that were falsely assigned among cells can be filtered out using noise detection in DPC^18^ or cell stains. which can be analyzed downstream for purposes such as cell typing, *etc*. (Fig. 1b (IV)). To illustrate this framework, we applied ClusterMap to representative *in situ* transcriptomics data from mouse brain tissue collected by the STARmap method^6^ (Fig. 1c).

Next, we examined and validated the performance of ClusterMap in diverse biological samples at different spatial scales in both 2D and 3D (Fig. 1d). First, based on the assumption that cellular RNAs have a different distribution in the nucleus or cytoplasm^19^, we used ClusterMap to cluster mRNAs within one cell to define the nuclear boundary. Here, RNA spots with both highly correlated neighboring composition and close spatial distances were merged into a single signature (Supplementary Fig. 1a, Methods). Then, a convex hull was constructed from the nucleus spots, denoting the nuclear boundary. The patterns of ClusterMap-constructed nuclear boundaries were highly correlated with DAPI stains, confirming the power of ClusterMap for segmentation at the subcellular resolution (Fig. 1d (I)). Second, we compared cell segmentation results by ClusterMap with conventional watershed^13^ segmentation (Methods) on the same mouse cortex cells. Comparing with the conventional watershed method, ClusterMap accurately identified cells, more precisely outlined cell boundary and illustrated cell morphology (Fig. 1d (II)). Last, we extended ClusterMap to diverse tissue types at different scales in both 2D and 3D, where dense heterogeneous populations of cells with arbitrary shapes exist. Cell identification results for the mouse cerebellum, the ileum, and the cortex are shown in Fig. 1d (III)-(V).

### Spatial clustering analysis in mouse brain

We first demonstrated ClusterMap on the mouse primary visual cortex from the STARmap mouse primary cortex (V1) 1020-gene dataset^6^ (Supplementary Table 1). When sequenced transcripts were more likely to populate the cytoplasm, sparsely sampled spots based on DAPI signals were combined with RNAs to compensate for the lack of signals in the centers of cells, and they were together processed with modified ClusterMap procedures (Fig. 2a, Methods). The results show clear cell segmentation even for strongly crowded mouse V1 cortex cells (Fig. 2b). Additionally, we evaluated whether ClusterMap-identified cell center coordinates were within corresponding expert-labeled cell regions on eight STARmap mouse V1 datasets to validate its accuracy (Supplementary Fig. 1b,c). Notably, ClusterMap cell labeling reached accuracy levels of 80-90% (Methods). Furthermore, we integrated gene expression in either ClusterMap-identified or expert-labeled cells with scRNA-seq data and compared their correlation^20,21^ (Supplementary Fig. 2d, Methods). Again, ClusterMap exhibited a comparable performance with expert-annotated segmentation. In the mouse V1 cortex dataset, ClusterMap identified cell types^22^ that matched both expression signature and tissue localization of the segmentation based on the previous report^8^ (Supplementary Fig. 2a-c). Importantly, ClusterMap can consistently identify cell types and their localization across biological replicates and in the mouse brain regions (Supplementary Fig. 2e-h).

**Fig. 2:**
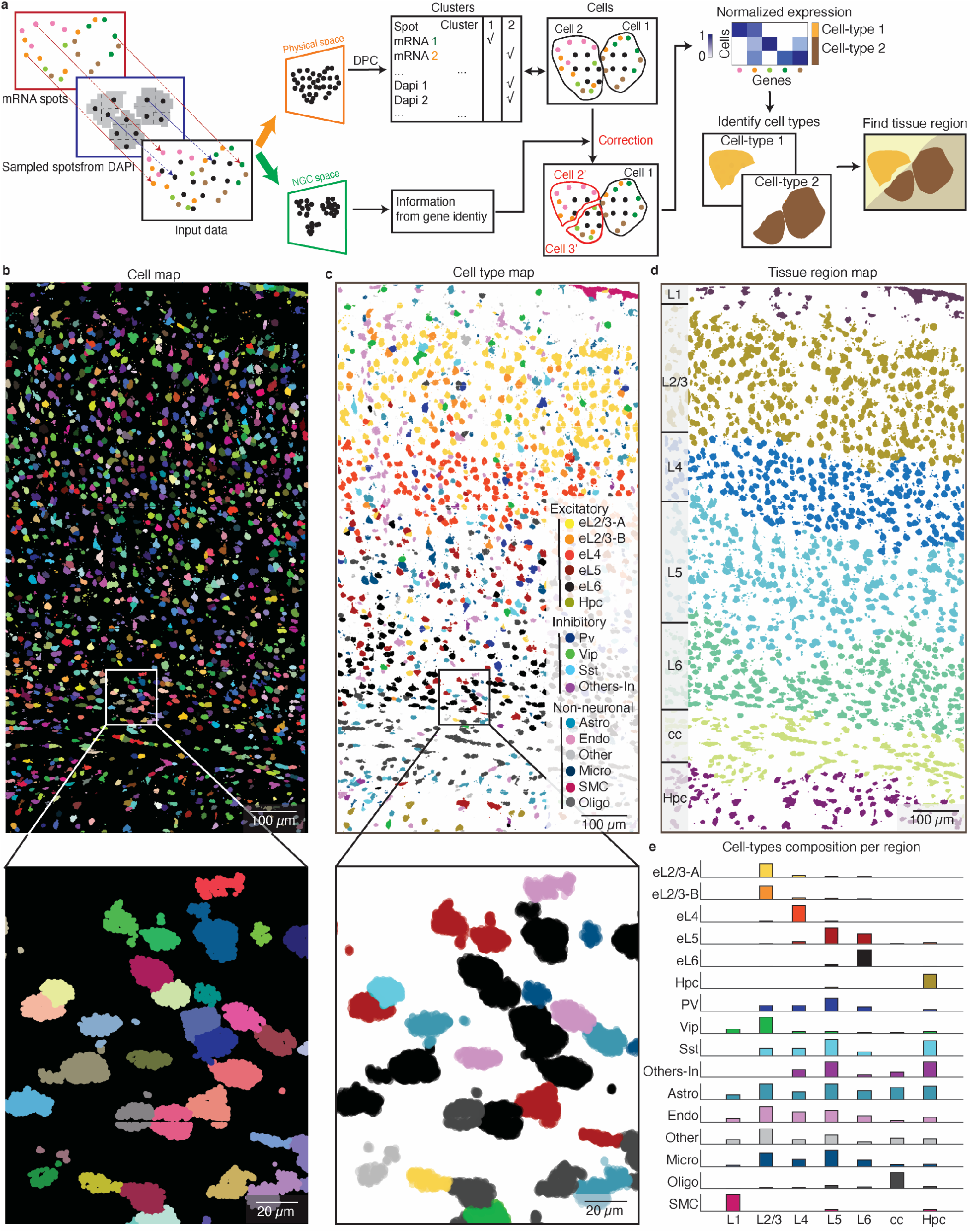
ClusterMap generates cell-type and tissue-region maps in mouse primary cortex (V1). **a**, Workflow of ClusterMap method that integrates DAPI signals for spatial clustering. **b**-**d**, ClusterMap generates cell (segmentation) map (**b**), cell-type map (**c**), and tissue region map (**d**) of the STARmap mouse V1 1020-gene dataset^6^, which includes 1,447 identified cells. **b**, mRNA molecules are color-coded by their cell attributes. **c**, The cell type names and colorings are from Ref (6). The number of cells in each cell type is as follows: eL2/3-A, 208; eL2/3-B, 42; eL4, 149; eL5, 118; eL6, 155; Pv, 37; Vip, 27; Sst, 40; Others-In, 18; Astro, 121; Endo, 134; SMC, 62; Micro, 150; Oligo-A, 164; Oligo-B, 12. Bottom panels in (**b, c**) show the zoomed-in views from the rectangular highlighted regions in upper panels. **d**, The tissue regions are segmented and cells in the same layer are shown in the same color. From top to bottom, the tissue region map shows: L1 to L6, the six neocortical layers; cc, corpus callosum; HPC, hippocampus. **e**, Bar plots of composition of 16 cell types across 7 layers. Values are normalized in each row. The colors correspond to the cell type legend in (**c**).

The next challenge was to apply ClusterMap on the cell-typing map to identify tissue regions. In this case, ClusterMap further clustered cells based on their physical and cell-type identity, providing similar clustering analyses of physical and high-dimensional cell-type information. ClusterMap computed a neighborhood cell-type composition (NCC) coordinates of each cell^23^ and then clustered joint physical and NCC coordinates of cells (Supplementary Fig. 1d, Methods). As a result, cells with both highly correlated neighboring cell-type composition and close spatial distances are clustered into a single tissue region signature. The results showed that ClusterMap accurately detected cortical layering, which allows for the quantification of cell-type composition of each cortical layer (Fig. 2b-e). The distinct region-specific distribution of excitatory neurons can be observed in the L2/3, L4, L5, and L6 canonical layers, while oligodendrocytes were significantly distributed within the corpus callosum layer. In summary, ClusterMap can effectively, accurately, and automatically conduct cell segmentation, cell typing, and tissue region identification.

### ClusterMap enables spatial clustering and cell-cell interaction analyses in mouse placenta

To further demonstrate the generality of ClusterMap, especially its applicability to tissues with high cell density and variable nuclear/cytosolic distribution of RNAs, we applied ClusterMap to the STARmap mouse placenta 903-gene dataset (Fig. 3a, b, Supplementary Table 1). With ClusterMap analyses described in Fig. 2a, up to 7,700 cells were identified (Supplementary Fig. 3a) and then clustered into eleven cell types using Louvain clustering^22^, which is consistent with cell types defined from scRNA-seq (Fig. 3e, f, Supplementary Fig. 3b-d). ClusterMap identified seven tissue regions based on the cell-type map (Fig. 3g, h). Further analysis showed that Regions IV and VI consisted of similar cell-type compositions, while region I consisted mostly of maternal decidua (MD) −2 cells.

**Fig. 3:**
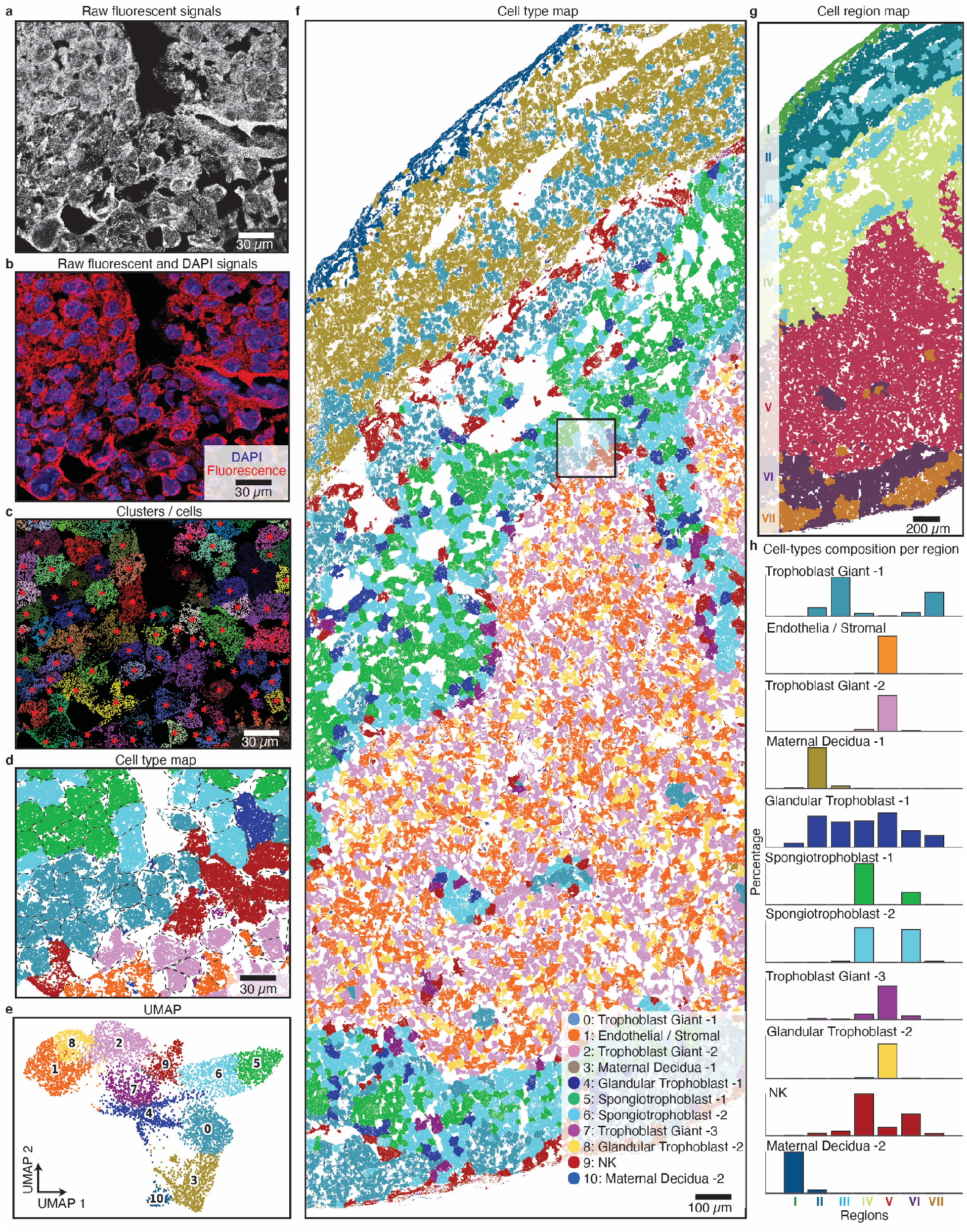
ClusterMap generates cell-type and tissue-region maps in mouse placenta. **a**, Raw fluorescent signals for a part in the STARmap mouse placenta 903-gene dataset^6^. Four-channel images in the first sequencing round are overlapped in grayscale to show the mRNA distribution. **b**, Composite image by overlapping (**a**) in red and DAPI signals in blue shows the distribution of mRNA relative to cell nuclei. A majority of mRNA molecules distributed outside the cell nucleus, resulting in holes in the cell center. **c, d**, ClusterMap generates cell map (**c**) and cell-type map (**d**) of (**a**). Panels (**a**-**d**) show the zoomed-in view from the highlighted rectangle in (**f**), the original dataset. **e**, Uniform manifold approximation plot (UMAP) shows clustering of 11 groups across 7,224 cells in the original placental dataset. **f**, Spatial organization of the cell types in the placental tissue section. The number of cells in each type is as follows: 0: Trophoblast Giant-1 (TG-1), 947; 1: Endothelial/Stromal (E/S), 924; 2: Trophoblast Giant-2 (TG-2), 921; 3: Maternal Decidua-1 (MD-1), 851; 4: Glandular Trophoblast-1 (GT-1), 706; 5: Spongiotrophoblast-1 (ST-1), 696; 6: Spongiotrophoblast-2 (ST-2), 680; 7: Trophoblast Giant-3 (TG-3), 550; 8: Glandular Trophoblast-2 (GT-2), 420; 9: NK, 392; 10: Maternal Decidua-2 (MD-2), 137. **g**, The spatial tissue region map of (**f**). **h**, Bar plots of composition of 11 cell types across 7 regions. Values are normalized in each row. Cell types in (**f, h**) are color-coded as in (**e**).

The discovery of the interwovenness of different tissue regions in placenta samples suggests the rich patterns of cell-cell interactions. We further sought to use ClusterMap results to characterize the near-range cell interaction networks by generating a mesh graph via Delaunay triangulation of cells and modeling the cellular relationships based on the i-niche concept^24^. In this way, we identified the nearest neighbors of each cell which were directly contacting each other (Fig. 4a-c) and quantified the average number of cells per cell-type among the first-tier neighbors (Fig. 4e), which could reveal crucial information about the affinity and communication among different cell types. Through this methodology, we discovered cell-type-specific cellular interactions: MD-1, MD-2, trophoblast giant-1 (TG-1), and NK cells mainly self-aggregate; glandular trophoblast-2 (GT-2), TG-2, TG-3, and endothelial/stromal cells widely connect with four types of cells; and Spongiotrophoblast −1 and Spongiotrophoblast −2 cells have a high affinity to each other. We envision that identifying cell-cell interaction facilitates future in-depth studies of the biological mechanisms and physiology of placenta and placenta-related diseases.

**Fig. 4:**
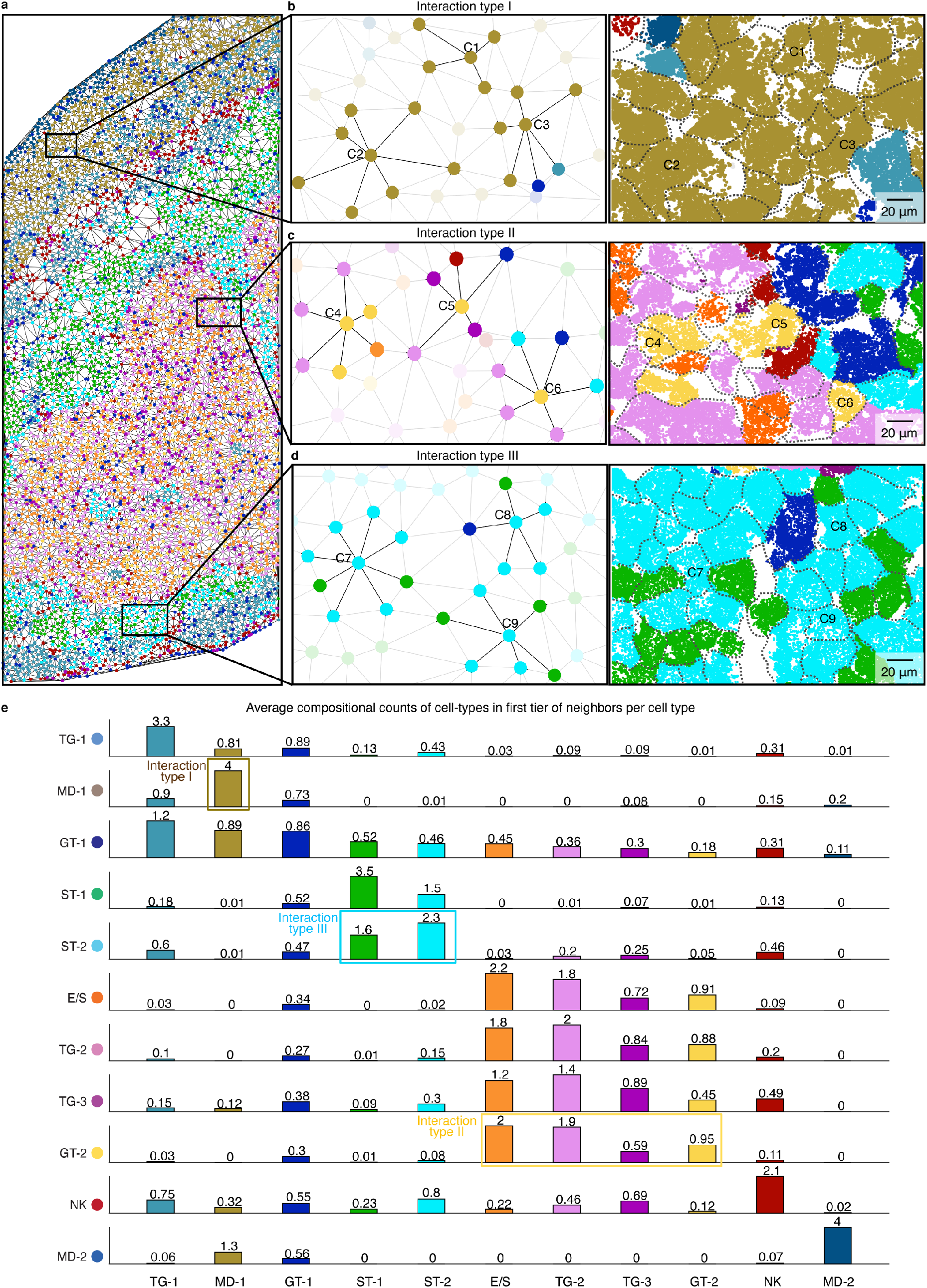
ClusterMap reveals cell-cell interactions in the placenta. **a**, Mesh graph generated by Delaunay triangulation^24^ of cells shown in the STARmap mouse placenta 903-gene reveals cell-cell interactions. Each cell is represented by a spot in the color of its corresponding cell type. Physically neighboring cells are connected via edges. **b**-**d**, A zoomed-in view of the top, middle, and bottom square in (**a**). The intercellular connection is centered on three MD-1 type (C1, C2, C3), GT-2 type (C4, C5, C6), and ST-2 (C7, C8, C9) type cells, respectively, with their first tier of neighboring cells highlighted. Left: schematic; right: cell map. **e**, Bar plots of the average number of cells per cell type among the first-tier neighbors, revealing clear patterns of cell-type specific cell-cell communication. Cells in Interaction Type I, II and III show selective association with cell types highlighted in the corresponding bounding box. The cell types on the axes are denoted by initialisms.

### ClusterMap is applicable across various *in situ* transcriptomic methods

Beyond just STARmap^6^, we further applied ClusterMap to analyze mouse brain tissue from three other *in situ* transcriptomics methods. Analyses of the imaged transcripts in the hypothalamic preoptic region by MERFISH^3^, the isocortex region by pciSeq^4^, and the somatosensory cortex by osmFISH^5^ are shown respectively in Fig. 5 (Supplementary Table 1). We used RNA spot matrices from subsets of the published data and applied ClusterMap analysis described in Fig.1b. Despite the differences in experimental designs and the number of transcript copies, ClusterMap identifies cells successfully. The identified cell types and their spatial patterns from ClusterMap were consistent with published results from conventional segmentation methods and scRNA-seq (Supplementary Fig. 4). Notably, ClusterMap can provide more detailed cell morphology and increase the number of identified cells (from 1,420 to 3,113 for MERFISH, from 893 to 1,962 for osmFISH).

**Fig. 5:**
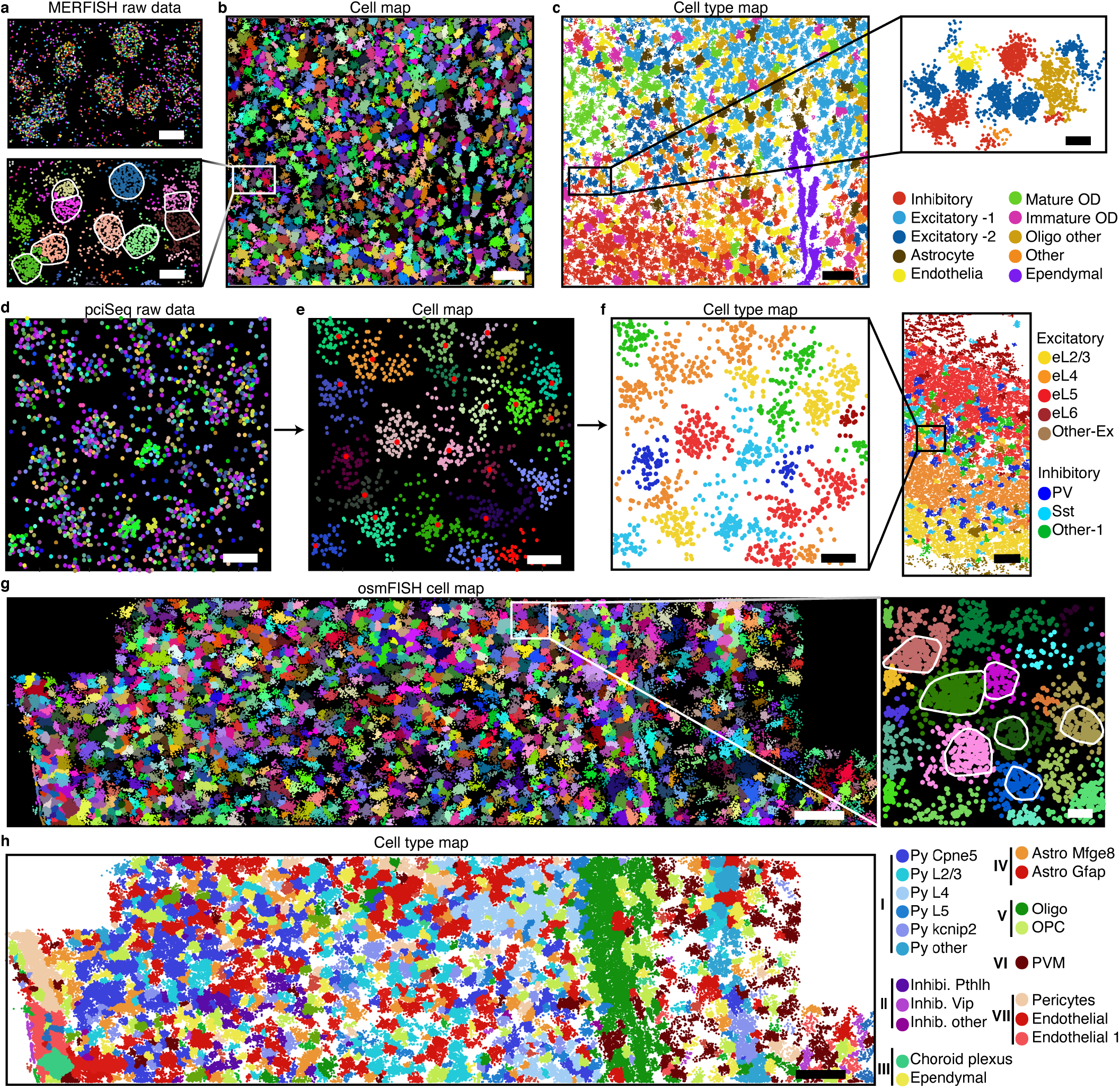
ClusterMap across various protocols. **a**, Raw spatial transcriptomics data from the MERFISH dataset^3^ (zoomed-in view corresponding to the highlighted rectangle in (**b**)). Different colors represent different gene types. Scale bar: 10 µm. **b**, ClusterMap generates the cell map of a selected area in MERFISH mouse POA dataset^3^, including 3,113 cells. Scale bar: 100 µm. Lower left: zoomed-in view of the highlighted square, the cell map of (**a**). Scale bar: 10 µm. White convex hulls are based on previous cell segmentation^3^. **c**, The spatial organization of cell types in (**b**). Scale bar: 100 µm. Upper right: zoomed in view of the highlighted rectangle, the cell type map of (**a**). Scale bar: 10 µm. **d**, Raw spatial transcriptomics data from the pciSeq mouse isocortex dataset^4^. **e**, ClusterMap generates the cell map of (**d**). Red points: the density peak of cells. Scale bar for (**d**,**e**): 5 µm. **f**, The cell type map in (**e**). Left: scale bar: 5 µm. Right: the spatial cell type map of the selected area in the pciSeq mouse isocortex dataset^4^. Scale bar: 100µm. **g**, The cell map of a part of the osmFISH mouse SSp dataset^5^, including 1,962 cells. Scale bar: 100µm. Left: zoomed-in view of the highlighted rectangle. White convex hulls are based on previously reported cell segmentation^5^. Scale bar: 10µm. **h**, The spatial organization of cell types in (**g**). Seven main types including I: Excitatory neurons; II: Inhibitory neurons; III: Ventricle; IV: Astro.; V: Oligodendrocytes; VI: Immune; VII: Vasculature^5^.

In conclusion, we analyzed mouse brain data from four representative *in situ* transcriptomic methods and validated the utility and universality of ClusterMap under different experimental methods. ClusterMap successfully produced comparable results across different methods with negligible modification applied.

### 3D ClusterMap analyses in thick tissue blocks

3D *in situ* transcriptomics data analysis is considered even more challenging because it is generally infeasible by manual labeling. However, 3D volumetric imaging and analysis are required to understand the structural and functional organization of complex organs. In this regard, exploring ClusterMap’s ability to analyze 3D *in situ* transcriptomics is particularly desired. We applied ClusterMap to two 3D thick-tissue samples: STARmap cardiac organoid 8-gene dataset^25^ and STARmap mouse V1 28-gene dataset^6^ (Supplementary Table 1). We analyzed the 3D data following the sample protocol described in Fig. 1b. In the 3D cardiac organoid sample, hierarchical clustering^26^ separated cells into three categories with distinct molecular signatures (Fig. 6a-c): *CD44* for mesenchymal stem cells (MSCs), *Nanog* for induced pluripotent stem cells (iPSCs) and four genes (*TNNI1, MYH7, MYL7, ATP2A2*) for cardiomyocytes (Supplementary Fig. 5a-c). The 100-μm-thick sample of mouse V1 includes all six cortical layers and the corpus callosum, in which up to 24,000 cells were identified and 3D clustered into eleven cell types (Fig. 6d, e, Supplementary Fig. 5d-g). Our results showed similar spatial distribution with previously published results, which used the conventional fluorescence image segmentation: excitatory neurons exhibited a gradient distribution, with the spatial density of each subtype gradually decaying to adjacent layers across the entire 3D space; inhibitory neurons showed a more dispersed distribution; and non-neuronal cells were largely located in the white matter and layer 1 (Fig. 6e). We can determine seven 3D tissue regions based on their corresponding cell-type compositions (Fig.6 f, g). We further characterized 3D cell-cell interactions in the mouse V1 and computed the average compositional neighboring cell types (Fig. 6h-k). In the minority inhibitory neurons, we observed a similar self-associative pattern as in previously published findings^6^: the nearest neighbor of any inhibitory neuron tends to be its own subtype. Three examples of inhibitory neuronal types (Pv, Sst, Vip) interactions are presented in Fig. 6h-j, respectively.

**Fig. 6:**
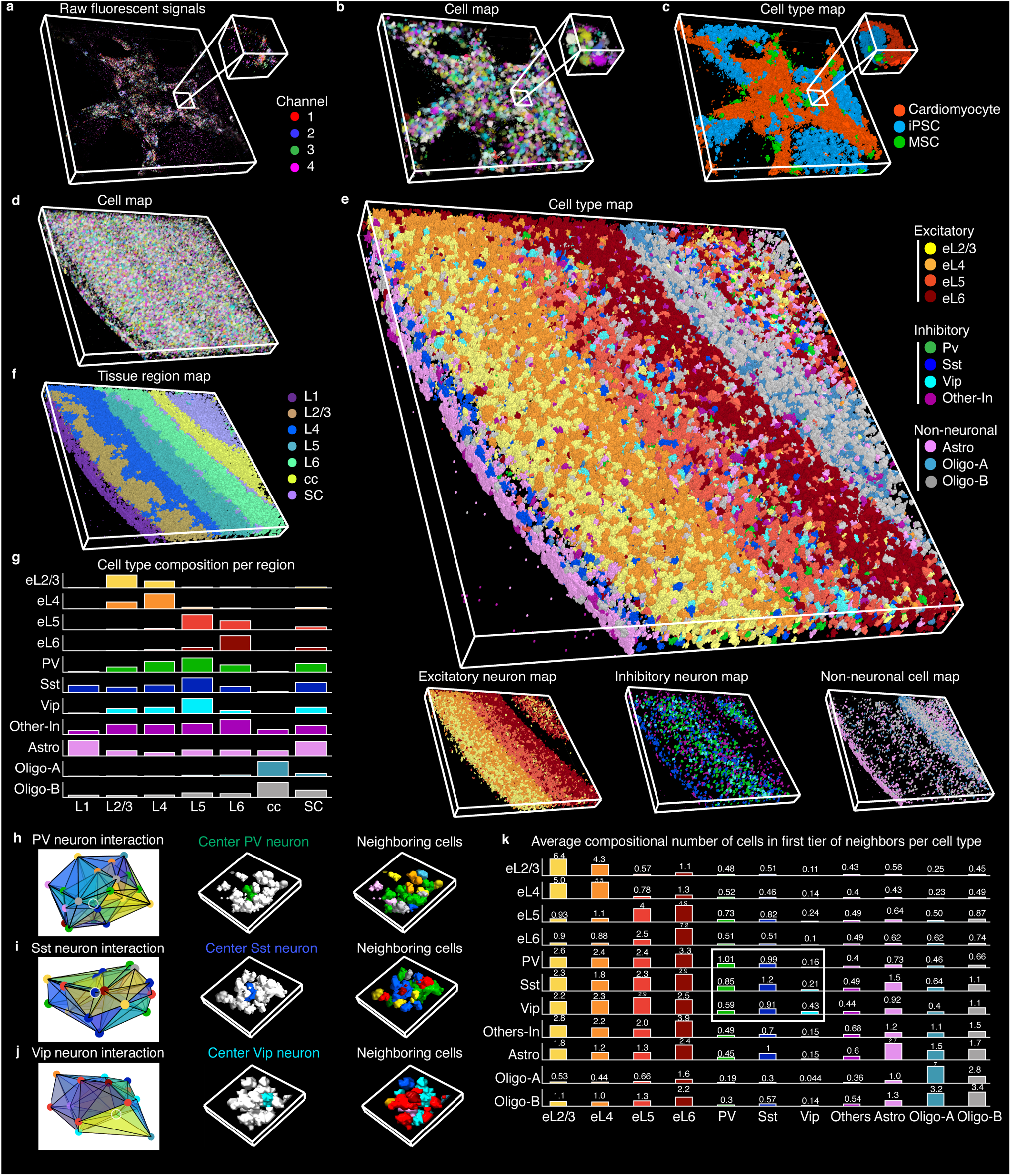
ClusterMap enables 3D *in situ* transcriptomics analysis. **a**, Raw fluorescent signals of 3D STARmap cardiac organoid 8-gene dataset. Width: 465 µm, height: 465 µm, depth: 97 µm. **b, c**, ClusterMap generates 3D cell map (**b**) and cell-type map (**c**) of (**a**), which includes 1,519 cells. Insets in (**a**-**c**) show zoomed-in views of the highlighted regions. **d**, ClusterMap generates a volumetric cell map of 3D STARmap mouse V1 28-gene dataset^6^, showing 24,590 cells. Width: 1545 µm, height: 1545 µm, depth: 100 µm. **e**, The 3D cell type maps of (**d**) show the spatial cell type distribution. **f**, The 3D tissue region map of (**e**). SC, subcortical. **g**, Bar plots of composition of 11 cell types across 7 tissue regions (layers). **h**-**j**, Example of cellular communication at a Pv, Sst, or Vip neuron, respectively. Left: schematics of 3D Delaunay triangulation of the Pv, Sst, or Vip neuron (highlighted in a white circle) and its first tier of neighboring cells. Middle: 3D spatial cell distribution of the first panel with the first tier of neighboring cells colored in white. Right: 3D spatial cell distribution of the first panel. Width 184 µm, height 194 µm, depth 100 µm. **k**, Bar plots of average composition of cell types around each cell type. Patterns of self-association in the minority inhibitory neurons are highlighted in the bounding box. Cell types in (**g**-**k**) are color-coded as in (**e**).

## Discussion

Spatial RNA localization intrinsically contains information related to biological structures and cell functions, which are yet to be effectively retrieved. ClusterMap exemplifies a computational framework that combines spatial and high-dimensional transcriptomic information from *in situ* single-cell transcriptomics to identify subcellular, cellular, and tissue structures in both 2D and 3D space. It is widely applicable to various experimental methods including, but not limited to, STARmap^6^, MERFISH^3^, pciSeq^4^, and osmFISH^5^. As a result, ClusterMap accurately created RNA-annotated subcellular and cellular atlases from *in situ* transcriptomic data across diverse tissue samples with different cell density, morphologies and connections, markedly expanded our knowledge of cellular organization across all scales from subcellular organelles through cell-type maps to organs, and enabled further characterization of the local microenvironment for individual cells. Our initial successful demonstration suggests that *in sit*u transcriptomic profiles contain unexplored biological and structural information that can be further extracted by new computational strategies.

Furthermore, ClusterMap is easy to scale up to a large dataset covering large-volume and organ-level imaging data. Beyond spatial transcriptomic data, ClusterMap can be generalized and applied to other 2D and 3D mapped high-dimensional discrete signals (*e*.*g*., proteins or signaling molecules imaging data)^27^. In the future, we envision that ClusterMap can also be extended by combining other types of biological features (*e*.*g*., subcellular organelles, cell shapes, *etc*.) to uncover the basic principles of how gene expression shapes cellular architecture and tissue morphology^28^.

## Methods

### Data pre-processing

#### 1. Thin-section STARmap Data Processing

All image processing steps were implemented using MATLAB R2019b and related open-source packages in Python 3.6 according to Wang *et al*., 2018^6^.

##### Image Preprocessing

For better unity of the illuminance and contrast level of the fluorescence raw image, a multi-dimensional histogram matching was performed on each image, which used the image of the first color channel in the first sequencing round as a reference.

##### Image Registration

Global image registration for aligning spatial position of all amplicons in each round of STARmap imaging was accomplished using a three-dimensional Fast Fourier transform (FFT) to compute the cross-correlation between two image volumes at all translational offsets. The position of the maximal correlation coefficient was identified and used to transform image volumes to compensate for the offset.

##### Spot Finding

After registration, individual spots were identified separately in each color channel on the first round of sequencing. For this experiment, spots of approximately 6 voxels in diameter were identified by finding local maxima in 3D. After identifying each spot, the dominant color for that spot across all four channels was determined on each round in a 5*5*3 voxel volume surrounding the spot location.

##### Spots and Barcode Filtering

Spots were first filtered based on fluorescence quality score. Fluorescence quality score is the ratio of targeted single-color channel to all color channels, which quantified the extent to which each spot on each sequencing round came from one color rather than a mixture of colors. Each spot is assigned with a barcode representing a specific kind of gene. The barcode codebook that contains all gene barcodes was converted into color space, based on the expected color sequence following 2-base encoding of the barcode DNA sequence^6^. Spot color sequences that passed the quality threshold and matched sequences in the codebook were kept and identified with the specific gene that that barcode represented; all other spots were rejected. The high-quality spots and associated gene identities in the codebook were then saved out for downstream analysis.

##### 2D Cell Segmentation

Two different methods were used to identify cell boundaries. First, the manually labeled segmentation masks from the original reference (Wang *et al*. 2018^6^) were obtained as baseline. Second, nuclei were automatically identified by the StarDist 2D machine learning model (Schmidt *et al*. 2018^15^) from a maximum intensity projection of the DAPI channel following the final round of sequencing. Then cell locations were extracted from the segmented DAPI image. Cell bodies were represented by the overlay of DAPI staining and merged amplicon images. Finally, a marker-based watershed transform was then applied to segment the thresholded cell bodies based on the combined thresholded cell body map and identified locations of nuclei. For each segmented cell region, a convex hull was constructed. Points overlapping each convex hull in 2D were then assigned to that cell, to compute a per-cell gene expression matrix.

#### 2. Thick-tissue STARmap Data Processing

##### 3D Image Registration

The displacement field of each imaging round was first acquired by registering the DAPI channel of each round to first-round globally by 3D FFT. Each sequencing image was applied with the corresponding transform of its round.

##### Spot Finding

After registration, individual spots were identified separately in each color channel on each round of sequencing. The extended local maxima in 3D were treated as an amplicon location. After identifying each spot, the dominant color for that spot across all four channels was determined on each round in a 3*3*3 voxel volume surrounding the spot location.

### Computation of Neighborhood Gene Composition (NGC)

To compute the neighborhood gene expression composition of each spot, we considered a spatially circular (2D) or spherical (3D) window over every spot (*S*) and counted the number of each gene-type among in the window. The raw count of each window was normalized to percentage for downstream analysis. The radius of the window *R* can be chosen either manually or by statistics to be close to the averaged size of organelles and cells for subcellular and single-cell analysis, respectively.

For a dataset with *T* kinds of sequenced gene, the definition of an NGC vector to the measured spot *i* is composed of the number of each gene-type windowed by radius *R* to the measured spot *i*.

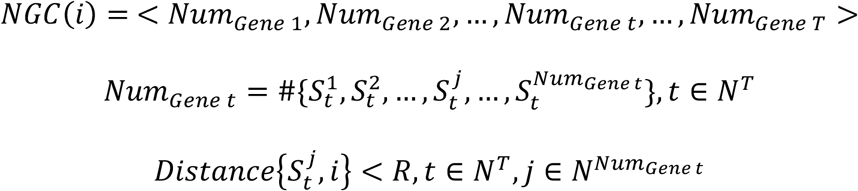

### Density peak clustering (DPC)

Based on the original DPC algorithm^18^, we first computed the two quantities: local density *ρ* and distance *δ* of every spot. We estimated the density by a Gaussian kernel with variance *d*_*c*_. The variance *d*_*c*_ is supposed to be close to the averaged radius *R* of cells for cellular segmentation. We can use *R* as *d*_*c*_. The definition of local density *ρ* and distance *δ* for spot *i* is:

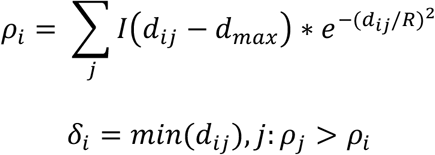

Note that *I*(*x*) = 1 *if x* < 0, *else I*(*x*) = 0, and *d*_*ij*_ is the distance between spot *i* and *j*. The optional parameter *d*_*max*_ is a striction on the maximum radius of the cell. For the point with the highest density, based on principles of DPC^18^, we took its distance value to the highest *δ* value. Note that for large data sets, the analysis is insensitive to the choice of *d*_*c*_ and results are robust and consistent.

After computing these two quantities of every spot, we generated a multiplication decision graph by computing *γ*, the product of *ρ* and *δ* and plotting every spot’s *γ* value in decreasing order.

Since the cell centers have both high local density and much higher distance at the same time, we chose the points with distinguishably higher *γ* values as cluster centers. We chose the ‘elbow point’ as the cutoff point in the multiplication decision graph where its *γ* value becomes no longer high and its change tends to be flat. *T* number of clusters *N* is equal to the number of points prior to the elbow point.

Next, we assigned each remaining point to one of the *N* clusters respectively in a descending order of *ρ* value in a single step manner. Each remaining spot was assigned to the same cluster as its nearest cluster-assigned neighbor. Each cluster was regarded as one cell. Finally, we filtered cells by limiting the minimum number of spots and genes expressed in one cell.

### Integration of the physical and NGC coordinate

The physical coordinates denote the spatial location of spots and the NGC coordinates denote the gene location of spots in a high-dimensional NGC space. For spot *i*, its physical and NGC coordinate are:

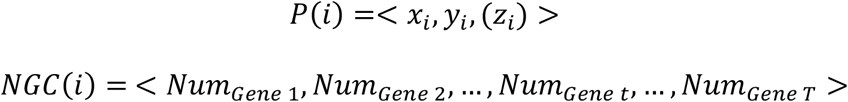

We used inversed Spearman correlation coefficient to measure the distance between two NGCs. Integration of these two coordinates can be distance-level, clustering-level, and guided-information based.

#### Distance-level integration

We computationally integrated the NGC and physical coordinates into the joint P-NGC coordinate over each spot. Specifically, we combined the physical and NGC distances information between *i* and its neighboring spots and used the joint distance as the metric to measure relationships between spots. Mathematically, the parameter *d*_*ij*_ used in the calculation of *ρ* and *δ* in DPC is:

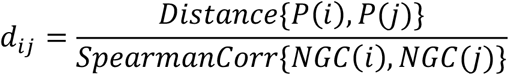

Then we performed the DPC algorithm and found the cells. This integration method is universal to any datasets. We used distance-level integration for MERFISH mouse POA^3^, pciSeq mouse isocortex^4^, osmFISH mouse SSp^5^, STARmap cardiac organoid 8-gene and STARmap mouse V1 28-gene dataset^6^.

#### Clustering-level integration

Since data points can be clustered by DPC using physical coordinates and NGC coordinates respectively, we can then do integration on the clustering level. To take these two variables into consideration, joint clustering methods can be explored. To take the correlations between variables into account, we can also optimize a pre-specified objective function. Here we don’t apply clustering-level integration to datasets presented in the manuscript.

#### Guided information-based integration

We first separated spots into clusters with physical coordinates and then corrected the clustering with guided information extracted from the NGC coordinates. To extract the guided information, we identified the neighbors of spot *i* that were at the distance of *R-2R* to spot *i* in the physical space. Then we computed these spots’ NGC distances to spot *i*. If the maximum of the NGC distances from spot *k* was higher than a threshold, we evaluated if spot *k* and spot *i* belong to the same cluster. If so, as they were both distant from spot *i* in physical and NGC spaces, this indicated the cell which spot *i* belongs to may be under-clustered. We counted the overall probability of each cell being missed and re-clustered the potentially incorrect cells with more than 50% missing probability. This integration method is best performed with datasets with DAPI stains. We used guided-information based integration for STARmap mouse V1 1020-gene^6^ and STARmap mouse placenta 903-gene dataset.

### Subcellular segmentation

To perform subcellular segmentation and construct nuclear boundaries we first computed the quantity NGC over each spot in an individual cell. The difference between NGC for subcellular segmentation and that for cellular segmentation is the radius of the window *R. R* should be either chosen manually or by statistics to be close to the averaged size of organelles. In addition, when the number of sequenced genes is limited, we can compute the NGC using a mesh graph by Delaunay triangulation of spots that models the relationship between RNA spots in the cell. A ring of spots that are neighbors of the central spot in the mesh graph is considered to locate most closely around the central cell. For a dataset with *TR* kinds of gene the definition of an NGC vector to the measured spot *i* is the composition of gene-types in its closest neighbors:

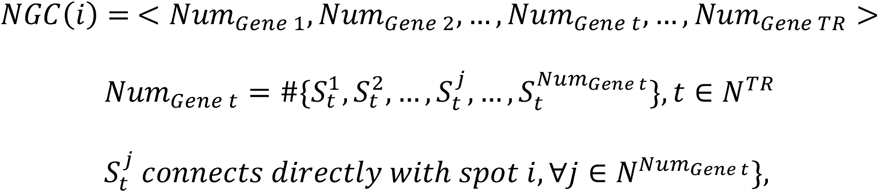

Then, similar to distance-level integration, we generate a joint P-NGC coordinate from the normalized NGC and physical coordinates over each spot:

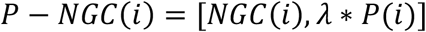

Here the optional parameter *λ* can control the influence of physical coordinates, depending on conditions. We then used K-means clustering to cluster spots into two regions with one for nucleus and one for cytoplasm. Finally, we constructed a convex hull based on the nucleus spots, denoting the nuclear boundary.

### Cell type classification

A two-level clustering strategy was applied to identify both major and sub-level cell types in the dataset. Processing steps in this section were implemented using Scanpy v1.6.0 and other customized scripts in Python 3.6 and applied according to Wang *et al*., 2018^6^. After filtration, normalization, and scaling, principal-components analysis (PCA) was applied to reduce the dimensionality of the cellular expression matrix. Based on the explained variance ratio, the top PCs were used to compute the neighborhood graph of observations. Then the Louvain algorithm^22^ was used to identify well-connected cells as clusters in a low dimensional representation of the transcriptomics profile. Clusters enriched for the excitatory neuron marker *Slc17a7* (vesicular glutamate transporter), inhibitory neuron marker *Gad1*, were manually merged to form two neuronal cell clusters, and then other cells represented non-neuronal cell populations. The cells were displayed using the uniform manifold approximation and projection (UMAP) and color-coded according to their cell types. The cells for each top-level cluster were then sub clustered using PCA decomposition followed by Louvain clustering^22^ to determine sub-level cell types.

### Construct tissue regions

#### 1. Neighborhood Cell-type Composition (NCC)

To construct tissue regions, we computed a global quantity: Neighborhood Cell-type Composition (NCC) over each cell (*C*). We considered a spatially circular (2D) or spherical (3D) window over every cell and estimated the composition of cell-types in the window. The radius of the window *RC* was chosen manually or by statistics of distances between cells to be as reasonable as possible.

For a dataset with *TC* kinds of gene, the definition of an NCC vector of the measured cell *i* was the composition of cell-types in the defined window that had radius *RC* to the measured cell *i*.

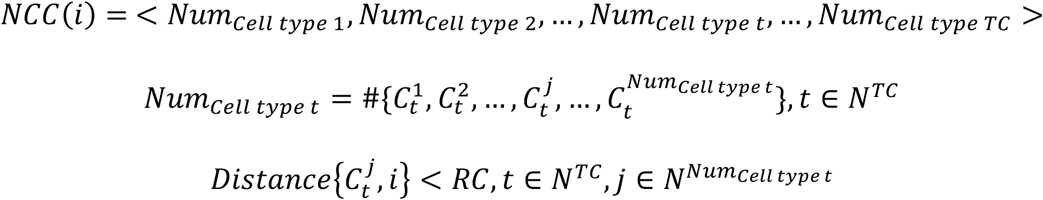

#### 2. K-means clustering

Tissue region signatures were identified using information from both NCC and physical locations of cells. Then we generated a joint P-NCC coordinate from normalized NCC and physical coordinates over each cell:

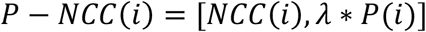

Here the optional parameter *λ* can control the influence of physical coordinates based on conditions. We then used K-means clustering on these high dimensional P-NCC coordinates to cluster cells into a pre-defined number of regions. Finally, we projected spatially back onto the cell-type map.

### Compare with expert-annotated labels

We evaluated the accuracy of cell identification by ClusterMap with corresponding eight expert annotated STARmap^6^ datasets (Supplementary Fig. 1c). Cells defined by ClusterMap consist of spots with physical locations while labels in the expert annotated STARmap datasets are connected components. We defined the accuracy as the percentage of ClusterMap-identified cells of which the center coordinates are correctly located within the labeled cells connected components. In other words, for each labeled connected component, we checked if there was only one cell center (cluster peak defined in DPC^18^) within the region. More than one cell centers were counted as over-clustering and no cell centers as under-clustering. The percentage was calculated by dividing the number of all incorrect cells (including over-clustered and under-clustered ones) by the total number of cells identified by ClusterMap.

### Integration with scRNA-seq

The cell identification performance was validated by performing a leave-one-out benchmark. Before integration^20,21^, the scRNA-seq and *in situ* sequencing data were preprocessed using the Seurat package.

1. Log-normalization: Divide the gene counts for each cell by the total counts for that cell and multiply by the scale. Factor = 10,000. Then perform natural-log transformation using *log1p*.

2. Scaling the data: Subtract the average expression for each gene and divide the centered gene expression profiles by their standard deviation.

For a shared gene list of scRNA-seq and *in situ* sequencing data with *n* genes, one non-repeating gene was left out in each round, and the rest *n-1* genes were used for integration with scRNA-seq data and then the prediction of the left-out gene’s expression profile. The integration and prediction steps were performed using *FindTransferAnchors* and *TransferData* functions in Seurat, which identified anchors between the reference (scRNA-seq) and query (*in situ* sequencing) dataset in reduced dimensions (reduction = ‘cca’) using mutual nearest neighbors and used these anchors to predict the left-out gene expression.

Next, the Pearson correlation of measured and predicted profile was calculated as the benchmark metrics. Finally, we compared the correlation between ClusterMap or manual annotation with scRNA-seq and gave quantitative analyses using violin plot, which showed the distribution of correlation for different annotation methods, and scatter plot, which represented the correlation values of these two methods for each gene.

### Label transfer

Cell type labels from scRNA-Seq dataset were projected onto spatially resolved cells from STARmap dataset by using the Seurat v3 integration method according to Stuart *et al*. 2019^20^. First, both datasets were preprocessed (normalization & scaling) and a subset of features (*e*.*g*., genes) exhibiting high variability was extracted. For STARmap dataset, all genes profiled were used whereas in scRNA-Seq dataset, the top 2,000 most variable genes identified by “*FindVariableFeatures*” function were used in downstream integration. Then “*FindTransferAnchors*” (reduction = “cca”) and Transfer Data functions were used to map the labels onto spatially resolved cells from the STARmap dataset. After label transferring, 6,672 out of 7,224 cells were observed with high-confidence cell type predictions (prediction score > 0.5), 8 out of 10 cell types labels were resolved.

## Data availability

The STARmap mouse V1 1020-gene and 160-gene sets^6^, MERFISH mouse POA set^3^, pciSeq mouse isocortex set^4^, osmFISH mouse SSp set^5^, and STARmap mouse V1 28-gene set^6^ are available as supplementary information file of the orginal manuscript. The data that support the findings of this study are available from the corresponding author upon reasonable request.

## Acknowledgements

We thank Prof. Xiaole Shirley Liu and Jane Salant for their helpful comments to the manuscript. Y.H. acknowledges the James Mills Peirce Fellowship from the Graduate School of Arts and Sciences of Harvard University. J.L. acknowledges the support from the William F. Milton Fund. X.W. acknowledges the support from Searle Scholars Program.

## Author contributions

X.W. and Y.H. conceived the idea. X.W., J.L., and Y.H. designed the research. Y.H. developed the framework, performed computational and data analyses, and prepared the manuscript. X.T. provided critical discussions during the whole development. J.H. provided preprocessing of raw data and the cell-typing pipeline. H.Z. performed the validation of the Placental Dataset using scRNA-seq. J.R., H.S., Z.L., Q.L., A.A. and J.S. provided *in situ* transcriptomic STARmap data. X.W., J.L., Y.H., J.H., X.T., K.C. and A.L. critically revised the manuscript. X.W. supervised the study.

## References

1. Stark, R., Grzelak, M. & Hadfield, J. RNA sequencing: the teenage years. Nat. Rev. Genet. 20, 631–656 (2019).

2. Crosetto, N., Bienko, M. & van Oudenaarden, A. Spatially resolved transcriptomics and beyond. Nat. Rev. Genet. 16, 57–66 (2015).

3. Moffitt, J. R. et al. Molecular, spatial, and functional single-cell profiling of the hypothalamic preoptic region. Science 362, eaau5324 (2018).

4. Qian, X. et al. Probabilistic cell typing enables fine mapping of closely related cell types in situ. Nat. Methods 17, 101–106 (2020).

5. Codeluppi, S. et al. Spatial organization of the somatosensory cortex revealed by osmFISH. Nat. Methods 15, 932–935 (2018).

6. Wang, X. et al. Three-dimensional intact-tissue sequencing of single-cell transcriptional states. Science 361, eaat 5691 (2018).

7. Eng, C.-H. L. et al. Transcriptome-scale super-resolved imaging in tissues by RNA seqFISH. Nature 568, 235 (2019).

8. Lee, J. H. et al. Fluorescent in situ sequencing (FISSEQ) of RNA for gene expression profiling in intact cells and tissues. Nat. Protoc. 10, 442–458 (2015).

9. Perkel, J. M. Starfish enterprise: finding RNA patterns in single cells. Nature 572, 549–551 (2019).

10. Kishi, J. Y. et al. SABER amplifies FISH: enhanced multiplexed imaging of RNA and DNA in cells and tissues. Nat. Methods 16, 533–544 (2019).

11. Thomas, R. M. & John, J. A review on cell detection and segmentation in microscopic images. In 2017 International Conference on Circuit, Power and Computing Technologies (ICCPCT), 1–5 (2017).

12. Moen, E. et al. Deep learning for cellular image analysis. Nat. Methods 16, 1233–1246 (2019).

13. Coelho, LP., Shariff, A. & Murphy, R. F. Nuclear segmentation in microscope cell images: a hand-segmented dataset and comparison of algorithms. In 2009 IEEE International Symposium on Biomedical Imaging: From Nano to Macro, 518–521 (2009).

14. Arganda-Carreras, I. et al. Trainable Weka Segmentation: a machine learning tool for microscopy pixel classification. Bioinformatics 33, 2424–2426 (2017).

15. Schmidt, U., Weigert, M., Broaddus, C. & Myers, G. Cell detection with star-convex polygons. In Medical Image Computing and Computer Assisted Intervention-MICCAI 2018 (eds. Frangi, A. F. et al.) 265–273 (Springer International Publishing, 2018).

16. Lein, E., Borm, L. E. & Linnarsson, S. The promise of spatial transcriptomics for neuroscience in the era of molecular cell typing. Science 358, 64–69 (2017).

17. Nitzan, M., Karaiskos, N., Friedman, N. & Rajewsky, N. Gene expression cartography. Nature 576, 132–137 (2019).

18. Rodriguez, A. & Laio, A. Clustering by fast search and find of density peaks. Science 344, 1492–1496 (2014).

19. Wang, G. et al. Spatial organization of the transcriptome in individual neurons. Preprint at https://www.biorxiv.org/content/10.1101/2020.12.07.414060v1 (2020).

20. Stuart, T. et al. Comprehensive integration of single-cell data. Cell 177, 1888–1902 (2019).

21. Abdelaal, T., Mourragui, S., Mahfouz, A. & Reinders, M. J. T. SpaGE: spatial gene enhancement using scRNA-seq. Nucleic Acids Res. 48, e107 (2020).

22. Blondel, V. D., Guillaume, J.-L., Lambiotte, R. & Lefebvre, E. Fast unfolding of communities in large networks. J. Stat. Mech. 10, P10008 (2008).

23. Park, J. et al. Segmentation-free inference of cell types from in situ transcriptomics data. Preprint at https://www.biorxiv.org/content/10.1101/800748v1 (2019).

24. Goltsev, Y. et al. Deep profiling of mouse splenic architecture with CODEX multiplexed imaging. Cell 174, 968–981 (2018).

25. Li, Q. et al. Cyborg organoids: implantation of nanoelectronics via organogenesis for tissue-wide electrophysiology. Nano Lett. 19, 5781–5789 (2019).

26. Rokach, L., Lior, R. & Oded, M. In Data Mining and Knowledge Discovery Handbook 321–352 (2005).

27. McCabe, A., Dolled-Filhart, M., Camp, R. L. & Rimm, D. L. Automated quantitative analysis (AQUA) of in situ protein expression, antibody concentration, and prognosis. J. Natl. Cancer Inst. 97, 1808–1815 (2005).

28. He, B. et al. Integrating spatial gene expression and breast tumor morphology via deep learning. Nat. Biomed. Eng. 666, 1–8 (2020).

## References

29. Bradski, G. The OpenCV library. Dr Dobb’s J. Software Tools 25, 120–125 (2000).

30. Goddard, T. D., Huang, C. C. & Ferrin, T. E. Visualizing density maps with UCSF Chimera. J. Struct. Biol. 157, 281–287 (2007).

31. Hunter, J. D. Matplotlib: a 2D graphics environment. Comput. Sci. Eng. 9, 90–95 (2007).

32. Jones, E., Oliphant, T. & Peterson, P. SciPy: open source scientific tools for Python. http://www.scipy.org/ (2001).

33. MacQueen, J. B. Some methods for classification and analysis of multivariate observations. In Proc. of the fifth Berkeley Symposium on Mathematical Statistics and Probability, 281–297 (University of California Press, Berkeley, 1967).

34. Higham, D. J. & Higham, N. J. MATLAB Guide, 150, (Siam, Philadelphia, 2016).

35. McInnes, L., Healy, J., & Melville, J. UMAP: uniform manifold approximation and projection for dimension reduction. Preprint at https://arxiv.org/abs/1802.03426 (2018).

36. McKinney, W. Data structures for statistical computing in Python. In Proc. 9th Python in Science Conference 51–56 (2010).

37. Oliphant, T. E. Guide to NumPy 1st edn 1, (Trelgol Publishing USA, 2006).

38. Pedregosa, F. et al. Scikit-learn: machine learning in Python. J. Machine Learn. Res. 12, 2825–2830 (2011).

39. Pérez, F., Granger, B. E. & Hunter, J. D. Python: an ecosystem for scientifc computing. Comput. Sci. Eng. 13, 13–21 (2011).

40. Rueden, C. T. et al. ImageJ2: ImageJ for the next generation of scientific image data. BMC Bioinformatics 18, 529 (2017).

41. Heideman, M., Johnson, D., & Burrus, C. Gauss and the history of the fast Fourier transform. IEEE ASSP Magazine 1, 14–21 (1984).

42. van derWalt, S. et al. scikit-image: image processing in Python. Peer J. 2, e453 (2014).

