## Supplementary information for "ClusterMap: multi-scale clustering analysis of spatial gene expression"

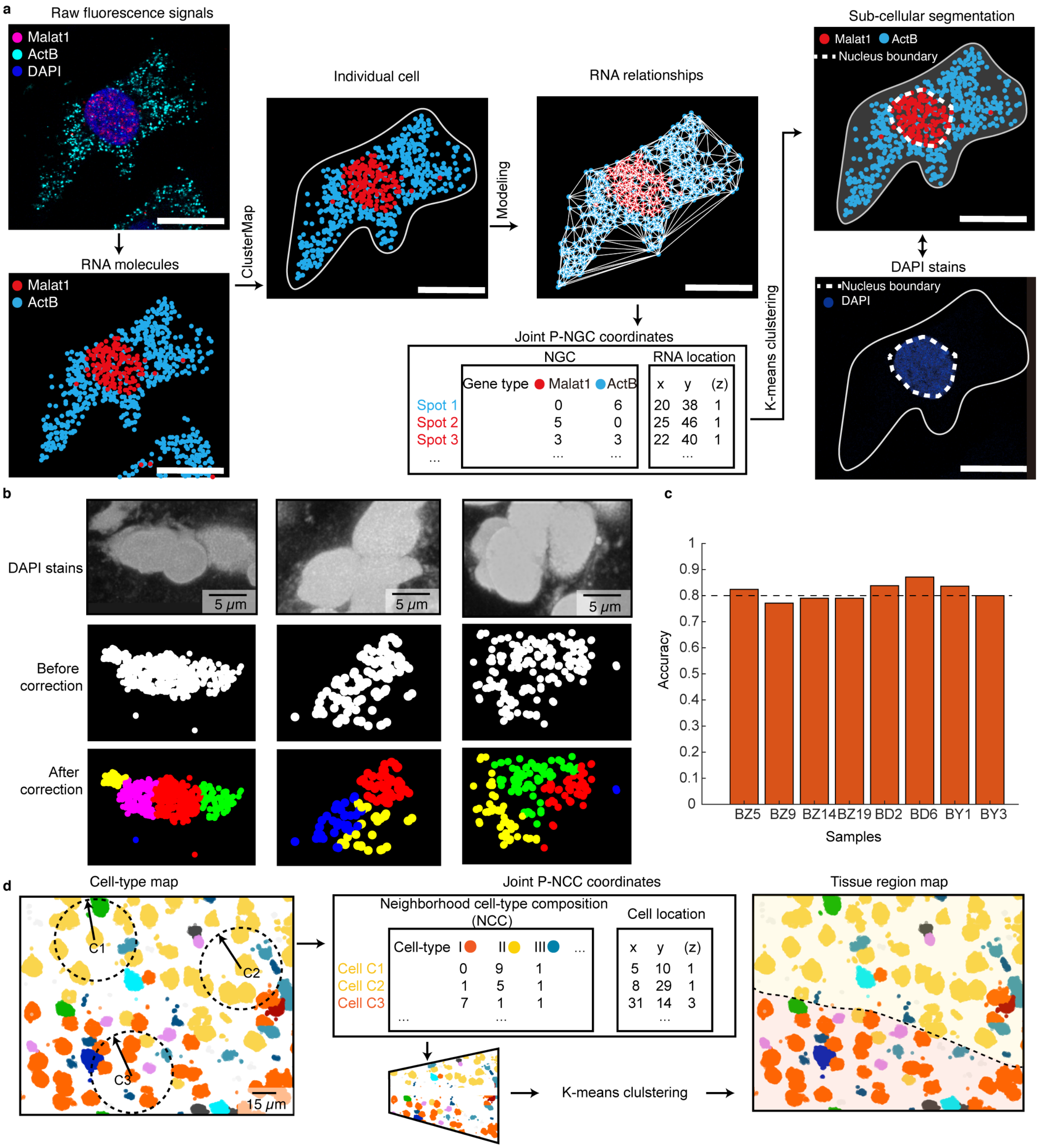
a
Raw fluorescence signals
Malat1
ActB
DAPI
RNA molecules
Malat1
ActB
ClusterMap
Individual cell
Modeling
RNA relationships
Joint P-NGC coordinates
NGC
RNA location
Gene type Malat1 ActB
x y (z)
Spot 1 0 6 20 38 1
Spot 2 5 0 25 46 1
Spot 3 3 3 22 40 1
K-means clulstering
Sub-cellular segmentation
Malat1 ActB
Nucleus boundary
DAPI stains
Nucleus boundary
DAPI
b
DAPI stains
5 μm
Before correction
After correction
c
Accuracy
Samples
BZ5 BZ9 BZ14 BZ19 BD2 BD6 BY1 BY3
0 0.1 0.2 0.3 0.4 0.5 0.6 0.7 0.8 0.9 1
d
Cell-type map
C1
C2
C3
15 μm
Joint P-NCC coordinates
Neighborhood cell-type composition (NCC)
Cell location
Cell-type I II III ...
x y (z)
Cell C1 0 9 1 5 10 1
Cell C2 1 5 1 8 29 1
Cell C3 7 1 1 31 14 3
K-means clulstering
Tissue region map

**Supplementary Figure 1**

Sub-cellular analysis, validation of the cell identification method, and tissue region analysis.

**a**, Subcellular analysis process the fourth panel IV in **Fig. 1d** by ClusterMap. A three-channel (magenta: *Malat1*; cyan: *ActB*; blue: DAPI) composite image shows raw fluorescent signals. After preprocessing mRNA molecules with specific genes are located, ClusterMap first performs cellular resolution and identifies individual cells. Then a mesh graph that models the relationships among mRNA spots in the cell is generated to compute the NGC coordinates and K-means clustering separate spots into two regions using joint physical and NGC coordinates. Finally a convex hull is constructed from the nucleus spots, denoting the nucleus boundary. The pattern of ClusterMap-constructed nucleus boundary is compared with the DAPI stains. Scale bar: 20μm.

**b**, Examples of the cell identification correction using information from NGC space during ClusterMap procedures in **Fig. 2a**. Upper: DAPI stains showing the cell nuclei. Middle: Cell clustering results using only information in the physical space. Closely overlapping cells are not separated. Lower: With information from NGC space, the under-clustered cells are separated.

**c**, The accuracy of cell identification results from eight STARmap datasets compared with corresponding expert-annotated labels. BZ5, BZ9, BZ14, BZ19: four STARmap 166-gene sets in mouse medial prefrontal cortex (mPFC); BD2, BD6: two STARmap 160-gene sets in mouse V1. BY1, BY3: two STARmap 1020-gene sets in mouse V1. The horizontal line is at 80% accuracy.

**d**, ClusterMap constructs the tissue regions after cell-typing. First, the neighborhood cell-type composition (NCC) of each cell is computed by considering a sliding window over the cell-type map. Then both the NCC and physical locations of cells are combined for K-means clustering.

Cells with highly correlated neighboring cell-type composition and close spatial distances are merged into a single tissue region signature.

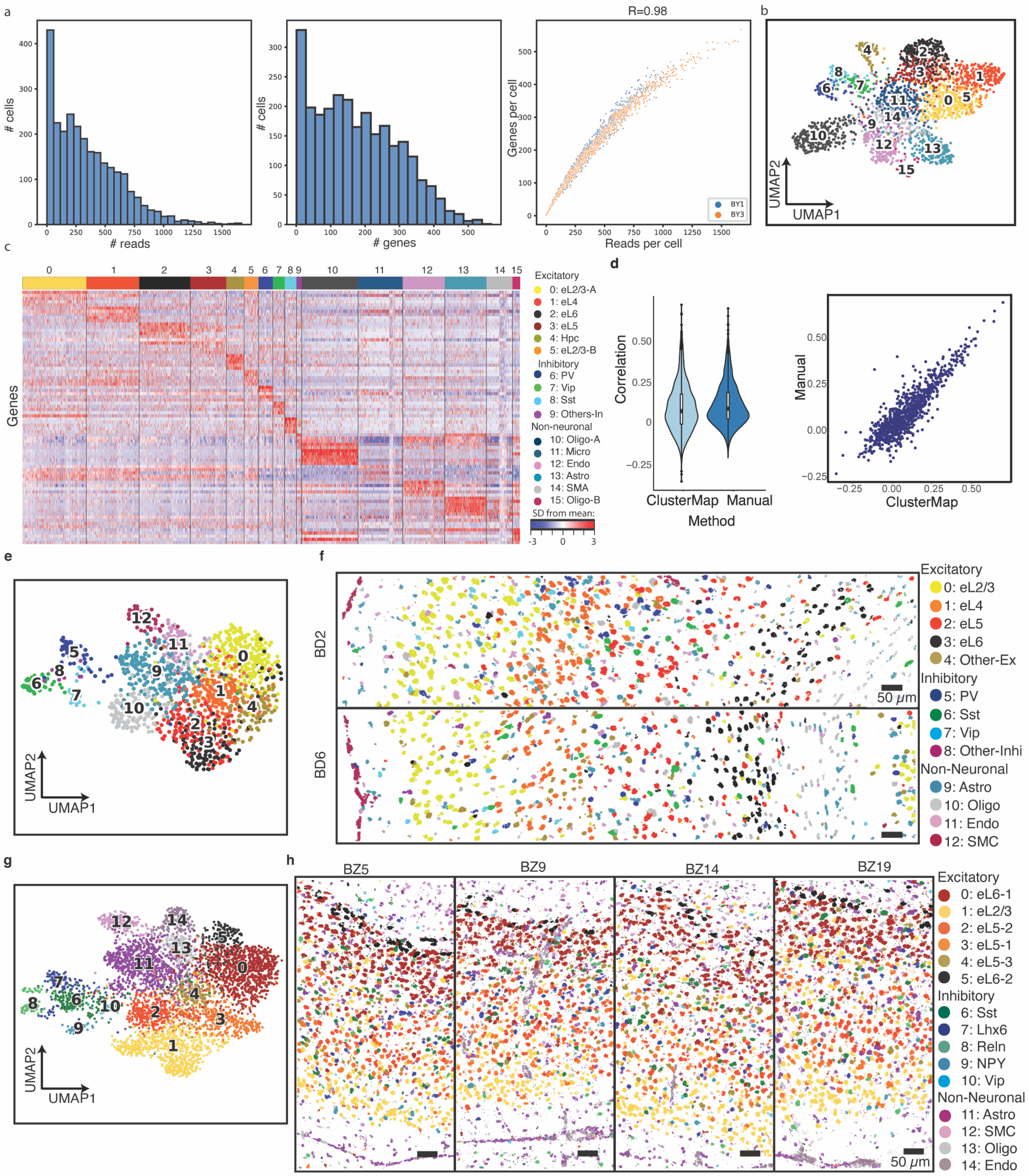

a
### cells
### reads
### genes
R=0.98
Genes per cell
Reads per cell
BY1
BY3
b
UMAP2
UMAP1
c
Genes
Excitatory
0: eL2/3-A
1: eL4
2: eL6
3: eL5
4: Hpc
5: eL2/3-B
Inhibitory
6: PV
7: Vip
8: Sst
9: Others-In
Non-neuronal
10: Oligo-A
11: Micro
12: Endo
13: Astro
14: SMA
15: Oligo-B
SD from mean:
d
Correlation
ClusterMap
Manual
Method
e
f
BD2
BD6
50 μm
Excitatory
0: eL2/3
1: eL4
2: eL5
3: eL6
4: Other-Ex
Inhibitory
5: PV
6: Sst
7: Vip
8: Other-Inhi
Non-Neuronal
9: Astro
10: Oligo
11: Endo
12: SMC
g
h
BZ5
BZ9
BZ14
BZ19
Excitatory
0: eL6-1
1: eL2/3
2: eL5-2
3: eL5-1
4: eL5-3
5: eL6-2
Inhibitory
6: Sst
7: Lhx6
8: Reln
9: NPY
10: Vip
Non-Neuronal
11: Astro
12: SMC
13: Oligo
14: Endo

**Supplementary Figure 2**

Identification of cell types in mouse V1 datasets.

**a**, Statistics of ClusterMap-identified cells in STARmap mouse V1 1020-gene (two replicates: BY1 and BY3). Left: Histogram of detected reads (DNA amplicons) per cell. Middle: Histogram of genes per cell. Right: Correlation plot between genes per cell and reads per cell.

**b**, **c**, UMAP and heatmap visualization of all excitatory, inhibitory and non-neuronal cell types in BY1 and BY3.

**d**, Correlation plots after integration with scRNA-seq atlas. Left: Violin plots of Pearson correlation between gene expression in scRNA-seq atlas and ClusterMap or manual. Right: Correlation plot between integration results of ClusterMap manual annotation.

**e**, **g**, UMAP visualization of all excitatory, inhibitory and non-neuronal cell types in STARmap 160-gene datasets in mouse V1 (two replicates: BD2, BD6, (**e**)), and STARmap 166-gene datasets in mPFC (four replicates, BZ5, BZ9, BZ14, BZ19, (**h**)).

**f**, **h**, Spatial organization map of cell types in BD2 and BD6 (**e**), and in BZ5, BZ9, BZ14 and BZ19 (**h**).

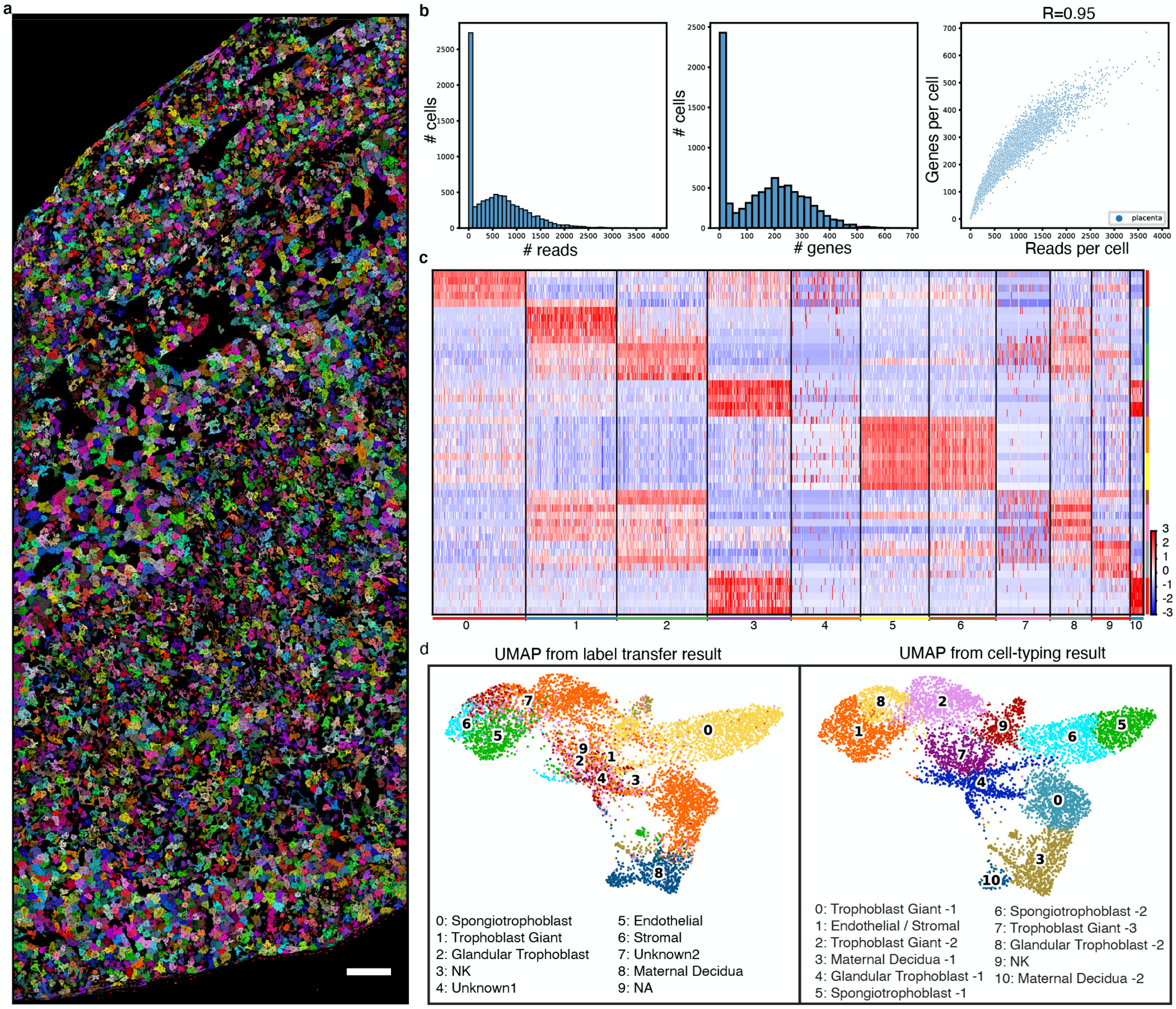

a
b
### cells
### reads
### genes
R=0.95
Genes per cell
Reads per cell
placenta
c
0
1
2
3
4
5
6
7
8
9
10
d
UMAP from label transfer result
UMAP from cell-typing result
0: Spongiotrophoblast
1: Trophoblast Giant
2: Glandular Trophoblast
3: NK
4: Unknown1
5: Endothelial
6: Stromal
7: Unknown2
8: Maternal Decidua
9: NA
0: Trophoblast Giant -1
1: Endothelial / Stromal
2: Trophoblast Giant -2
3: Maternal Decidua -1
4: Glandular Trophoblast -1
5: Spongiotrophoblast -1
6: Spongiotrophoblast -2
7: Trophoblast Giant -3
8: Glandular Trophoblast -2
9: NK
10: Maternal Decidua -2

**Supplementary Figure 3**

Analyses of the placental dataset.

**a**, ClusterMap generates the cell map of the STARmap mouse placenta 903-gene dataset, including 7,224 cells. Scale bar: 100 µm.

**b**, Statistics of ClusterMap identified placental cells as shown in (**a**). Left: Histogram of detected reads (DNA amplicons) per cell. Middle: Histogram of genes per cell. Right: Correlation plot between genes per cell and reads per cell.

**c**, Heatmap visualization of 11 cell types. Names are in the right panel of (**d**).

**d**, UMAP from label transfer results with scRNA-seq, compared with UMAP of the Louvain clustering in ClusterMap.

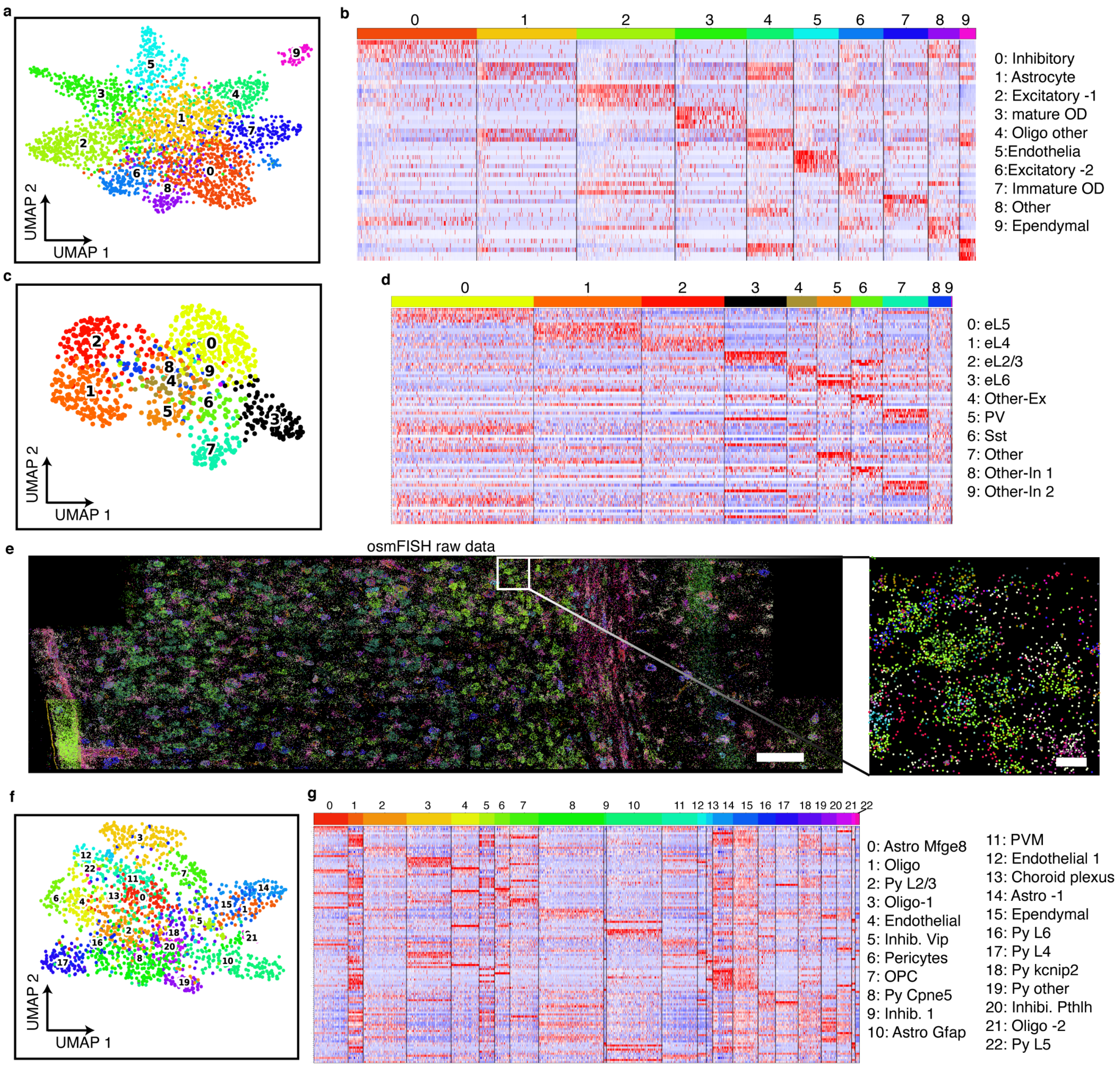
a
UMAP 2
UMAP 1
b
0: Inhibitory
1: Astrocyte
2: Excitatory -1
3: mature OD
4: Oligo other
5:Endothelia
6:Excitatory -2
7: Immature OD
8: Other
9: Ependymal
c
UMAP 2
UMAP 1
d
0: eL5
1: eL4
2: eL2/3
3: eL6
4: Other-Ex
5: PV
6: Sst
7: Other
8: Other-In 1
9: Other-In 2
e
osmFISH raw data
f
UMAP 2
UMAP 1
g
0: Astro Mfge8
1: Oligo
2: Py L2/3
3: Oligo-1
4: Endothelial
5: Inhib. Vip
6: Pericytes
7: OPC
8: Py Cpne5
9: Inhib. 1
10: Astro Gfap
11: PVM
12: Endothelial 1
13: Choroid plexus
14: Astro -1
15: Ependymal
16: Py L6
17: Py L4
18: Py kcnip2
19: Py other
20: Inhibi. Pthlh
21: Oligo -2
22: Py L5

**Supplementary Figure 4**

Analyses of datasets generated across various protocols.

**a**, **b**, UMAP and heatmap visualization of ten cell types in the selected area from the MERFISH mouse POA dataset.

**c**, **d**, UMAP and heatmap visualization of ten cell types in the selected area from the pciSeq mouse isocortex dataset.

**e**, Raw spatial transcriptomics data of the selected area from the osmFISH mouse SSp dataset. Scale bar: 100µm. Left: zoomed in view of the highlighted square. Scale bar: 10 µm.

**f**, **g**, UMAP and heatmap visualization of seven main types and 22 subtypes of (**e**).

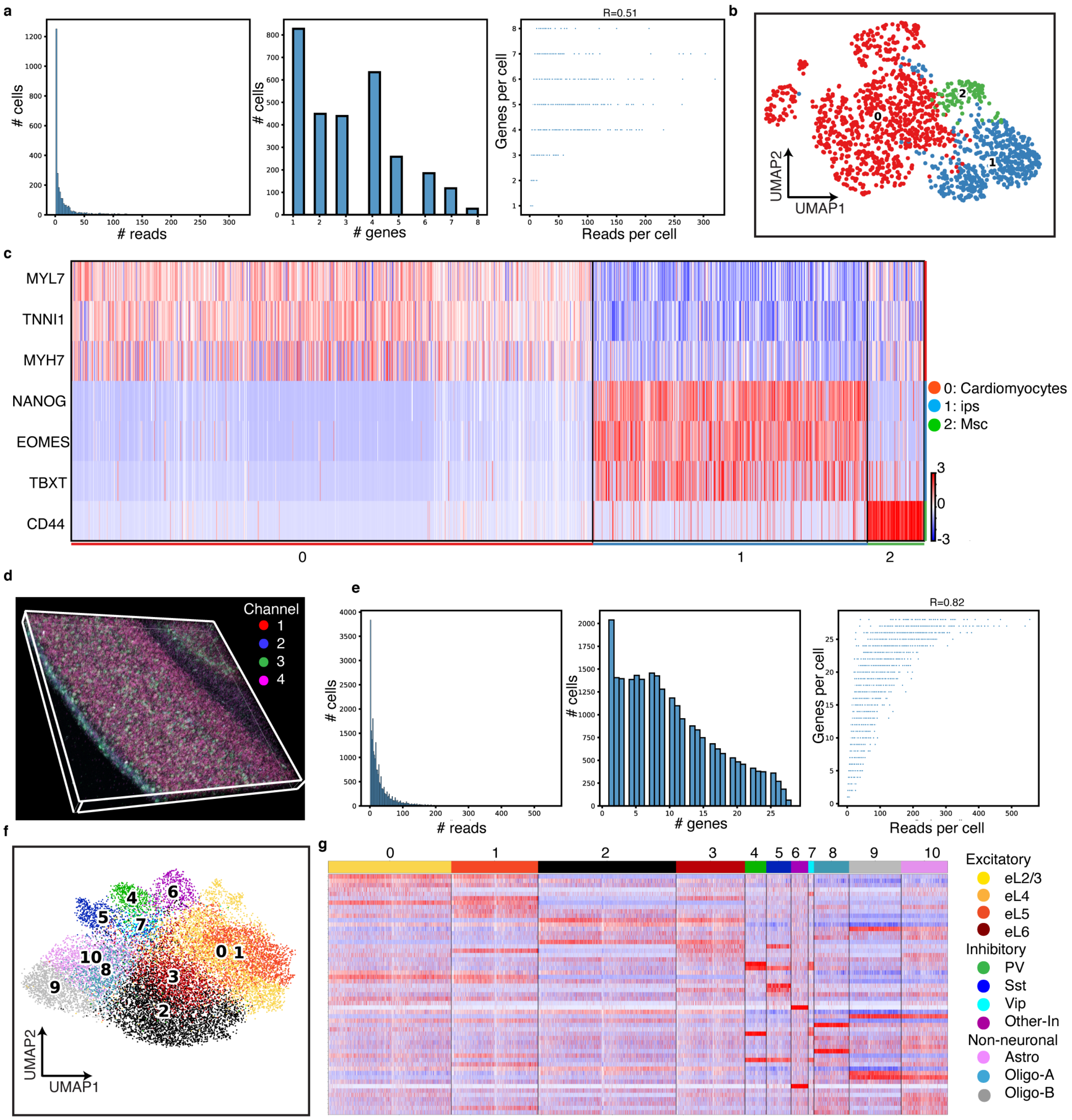
a
### cells
### reads
### genes
R=0.51
Genes per cell
Reads per cell
b
UMAP2
UMAP1
c
MYL7
TNNI1
MYH7
NANOG
EOMES
TBXT
CD44
0: Cardiomyocytes
1: ips
2: Msc
d
Channel
1
2
3
4
e
R=0.82
f
g
Excitatory
eL2/3
eL4
eL5
eL6
Inhibitory
PV
Sst
Vip
Other-In
Non-neuronal
Astro
Oligo-A
Oligo-B

**Supplementary Figure 5**

Analyses in the 3D datasets.

**a**, Statistics of ClusterMap identified cells in the 3D STARmap cardiac organoid 8-gene dataset. Left: Histogram of detected reads (DNA amplicons) per cell. Middle: Histogram of genes per cell. Right: Correlation plot between genes per cell and reads per cell.

**b**, **c**, UMAP and heatmap visualization of three cell types in the STARmap cardiac organoid 8-gene dataset. The number of cells in each cell type is as follows: cardiomyocytes, 929; induced pluripotent stem cells (iPSCs), 489; mesenchymal stem cells (MSCs), 101.

**d**, 3D four-channel composite raw fluorescent image of the first sequencing round shows spatial arrangement of mRNA molecules in the STARmap mouse V1 28-gene dataset. Width 184 µm, height 194 µm, depth 100 µm.

**e**, Statistics of ClusterMap identified cells in (**d**). Left: Histogram of detected reads (DNA amplicons) per cell. Middle: Histogram of genes per cell. Right: Correlation plot between genes per cell and reads per cell.

**f**, **g**, UMAP and heatmap visualization of three cell types of (**d**).

#### Supplementary Tables

##### Supplementary Table 1

| Dataset | Method | Tissue | # Gene | # Cell | # Cell type | Figure | Notes |
|---|---|---|---|---|---|---|---|
| STARmap mouse V1 1020-gene | STARmap | Mouse brain primary visual cortex | 1,020 | 1,447 | 16 | Fig. 1c, Fig. 2, Supplementary Fig. 2 | Source: Ref. 6. 2D analysis. |
| STARmap mouse placenta 903-gene | STARmap | Mouse brain primary visual cortex | 903 | 7224 | 11 | Fig. 3, Fig. 4, Supplementary Fig. 3 | New data. 2D analysis. |
| MERFISH mouse POA | MERFISH | Mouse brain hypothalamic preoptic region | 140 | 3,113 | 10 | Fig.5, Supplementary Fig. 4 | Source: Ref. 3. 2D analysis. |
| pciSeq mouse isocortex | pciSeq | Mouse brain isocortex region | 98 | 982 | 8 | Fig.5, Supplementary Fig. 4 | Source: Ref. 4. 2D analysis. |
| osmFISH mouse SSp | osmFISH | Mouse brain somatosensory cortex | 33 | 1,962 | 19 | Fig.5, Supplementary Fig. 4 | Source: Ref. 5. 2D analysis. |
| STARmap cardiac organoid 8-gene | STARmap | Cardiac organoid | 8 | 1,519 | 3 | Fig. 6, Supplementary Fig. 5 | New data, 3D analysis. |
| STARmap mouse V1 28-gene | STARmap | Mouse brain primary visual cortex | 28 | 24,590 | 11 | Fig. 6, Supplementary Fig. 5 | Source: Ref. 6. 3D analysis. |

Summary of the name, *in situ* sequencing protocol, tissue, number of genes, number of cells, number of cell types, corresponding figures and note of 7 datasets.
